# Dynein light chain-dependent dimerization of Egalitarian is essential for maintaining oocyte fate in *Drosophila*

**DOI:** 10.1101/2021.05.12.443861

**Authors:** Hannah Neiswender, Chandler H. Goldman, Rajalakshmi Veeranan-Karmegam, Graydon B. Gonsalvez

## Abstract

Egalitarian (Egl) is an RNA adaptor for the Dynein motor and is thought to link numerous, perhaps hundreds, of mRNAs with Dynein. Dynein, in turn, is responsible for the transport and localization of these mRNAs. Studies have shown that efficient mRNA binding by Egl requires the protein to dimerize. We recently demonstrated that Dynein light chain (Dlc) is responsible for facilitating the dimerization of Egl. Mutations in Egl that fail to interact with Dlc do not dimerize, and as such, are defective for mRNA binding. Consequently, this mutant does not efficiently associate with BicaudalD (BicD), the factor responsible for linking the Egl/mRNA complex with Dynein. In this report, we tested whether artificially dimerizing this Dlc-binding mutant using a leucine zipper would restore mRNA binding and rescue mutant phenotypes in vivo. Interestingly, we found that although artificial dimerization of Egl restored BicD binding, it only partially restored mRNA binding. As a result, Egl-dependent phenotypes, such as oocyte specification and mRNA localization, were only partially rescued. We hypothesize that Dlc-mediated dimerization of Egl results in a three-dimensional conformation of the Egl dimer that is best suited for mRNA binding. Although the leucine zipper restores Egl dimerization, it likely does not enable Egl to assemble into the conformation required for maximal mRNA binding activity.

## INTRODUCTION

Numerous mRNAs are specifically localized within the cytoplasm of *Drosophila* oocytes and embryos (Suter, 2018). In fact, establishment of oocyte fate, formation of Anterior-Posterior and Dorsal-Ventral polarity, and development of the embryo, all rely on correct localization of specific mRNAs (Weil, 2014). Although several mechanisms can contribute to the localization of mRNA, in many instance, localization requires the minus-end directed microtubule motor, cytoplasmic Dynein (hereafter Dynein) (Goldman and Gonsalvez, 2017). The primary RNA binding protein tasked with linking these diverse transcripts with Dynein is Egalitarian (Egl) (Dienstbier et al., 2009).

Dienstbier and colleagues demonstrated several years back that Egl is able to specifically interact with localization elements found in numerous Dynein-localized cargoes (Dienstbier et al., 2009). In addition to binding RNA, Egl also interacts with BicaudalD (BicD) and Dynein light chain (Dlc/Lc8) (Mach and Lehmann, 1997; Navarro et al., 2004). BicD is a highly conserved adaptor of Dynein and directly interacts with components of the Dynein motor and also with Dynactin, a regulator of Dynein activity (Hoogenraad and Akhmanova, 2016). Dlc is a core component of the Dynein motor, but also appears to have Dynein-independent function. Most notably, recent studies indicate that Dlc can aid in the dimerization of proteins, particularly those such as Egl which harbor long stretches of disordered regions (Barbar, 2008; Rapali et al., 2011; Reardon et al., 2020).

How does Egl link cargo with Dynein? Answering this questions has been the focus of several recent studies. The labs of Simon Bullock and Kathleen Trybus elegantly demonstrated that once Egl binds to cargo, its interaction with BicD is greatly enhanced. This cargo bound complex then associates with Dynein and is required for fully activating Dynein-mediated transport (McClintock et al., 2018; Sladewski et al., 2018). Our lab recently determined that Dlc is required for Egl dimerization (Goldman et al., 2019). Mutations in Egl that compromised Dlc binding were defective for dimerization. These mutants also displayed reduced RNA binding activity, and as a consequence, BicD binding was reduced (Goldman et al., 2019).

The current model in the field is that Egl dimerizes with the aid of Dlc. Dimeric Egl is then able to associate with mRNA cargo. This complex now has a high affinity for BicD, which in turn tethers the entire assemblage onto Dynein for processive minus-end microtubule transport (illustrated in Fig. 1). In support of this model, we showed that artificially dimerizing a mutant version of Egl that could not associate with Dlc was sufficient to restore its ability to bind to mRNA and BicD. These experiments were conducted using lysates from *Drosophila* S2 cells (Goldman et al., 2019). An important question raised by this finding is whether artificially dimerizing Egl could rescue mutant phenotypes in vivo. Answering this question has been the focus of the present study. Our results indicate that artificially dimerizing Egl does in fact restores BicD binding. However, in vivo, mRNA binding is only partially restored. This partial rescue results in inefficient mRNA localization, particularly at early stages of oogenesis. We posit that the artificially dimerized Egl assembles into a conformation that does not completely mimic Dlc-mediated dimeric Egl. As such, its capacity to bind mRNA is not fully restored. Thus, Dlc performs an essential role in facilitating Egl’s RNA binding activity. Disrupting this function results in mRNA localization defects and a failure to maintain oocyte fate.

**Figure 1:**
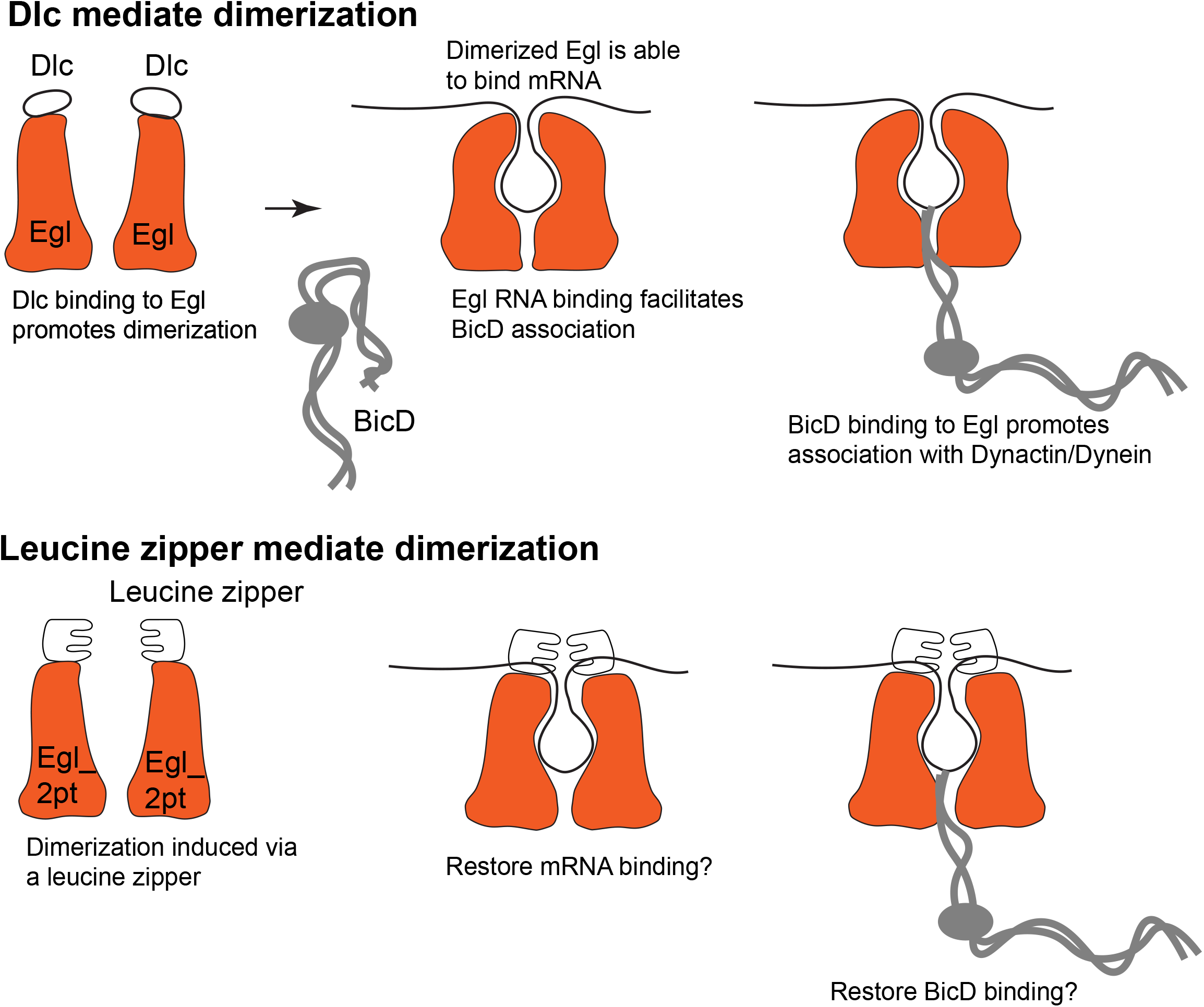
(Top panel) An illustration representing the current model. In the native context, Egl dimerization is facilitated by Dlc. Once dimerized, Egl is able to bind more efficiently with mRNA cargo. This complex now has a higher affinity for the Dynein adaptor, BicD. This binding relieves the auto-inhibition of BicD and the complex can now be linked with Dynein. (Bottom panel) In this report, we test whether artificially dimerizing the Egl_2pt mutant using a leucine zipper is able to restore mRNA binding, BicD binding and in vivo phenotypes.

## RESULTS

### Artificial dimerization of Egl restores BicD binding and partially restores mRNA binding

We recently demonstrated that Dlc is required for dimerization of Egl (Goldman et al., 2019). An Egl mutant that was unable to interact with Dlc (*egl_2pt*) was not only dimerization defective, it was also compromised for binding mRNA and BicD (Goldman et al., 2019; Navarro et al., 2004). In order to test the hypothesis that Egl dimerization is required for mRNA binding, and consequently associating with BicD, we attached a leucine zipper, a well-characterized dimerization domain, to the C-terminus of *egl_2pt*. The leucine zipper used in these studies was from the yeast GCN4 protein (Abel and Maniatis, 1989; Hope and Struhl, 1987). This mutant was expressed in S2 cells and tested for mRNA and BicD binding. As expected, Egl_2pt-zip did not bind Dlc, yet due to the presence of the leucine zipper, dimerization was restored (Goldman et al., 2019). Consistent with our hypothesis, using this in vitro assay, mRNA and BicD binding were also restored (Goldman et al., 2019) (illustrated in Fig. 1).

Our next objective was to determine whether artificial dimerization could restore Egl function in vivo. Transgenic flies expressing either FLAG or GFP tagged Egl_wt, Egl_wt-zip, Egl_2pt or Egl_2pt-zip were generated. The constructs contained silent mutations that made them refractory to depletion via *egl* shRNA (Sanghavi et al., 2016). This enabled us to generate flies that were depleted of endogenous Egl, yet expressed the transgenic constructs in the germline. For these initial binding experiments, an early-stage Gal4 driver (maternal alpha tubulin-Gal4; Bloomington stock 7063) was used to deplete endogenous Egl. This driver is not active in the germarium but is turned on in early stage egg chambers (Sanghavi et al., 2016). As such, the function of Egl in oocyte specification can be bypassed and mature egg chambers can be generated (Sanghavi et al., 2016), thus enabling us to perform the binding experiments.

We first tested the ability of these constructs to bind BicD and Dlc using a co-immunoprecipitation assay. Ovarian lysates were prepared from flies expressing GFP tagged Egl_wt, Egl_2pt, Egl_2pt-zip or Egl_wt-zip. The tagged proteins were immunoprecipitated using GFP-trap beads and the co-precipitating BicD and Dlc were analyzed using their corresponding antibodies. As expected, Egl_2pt displayed reduced binding to both Dlc and BicD (Fig. 2A, B). Although Egl_2pt-zip was still unable to bind to Dlc, its ability to associate with BicD was restored (Fig. 2A, B). Addition of the zipper to wild-type Egl did not enhance or diminish its ability to bind either BicD or Dlc (Fig. 2A, B). Thus, this approach revealed that artificial dimerization of Egl_2pt restores BicD binding.

**Figure 2:**
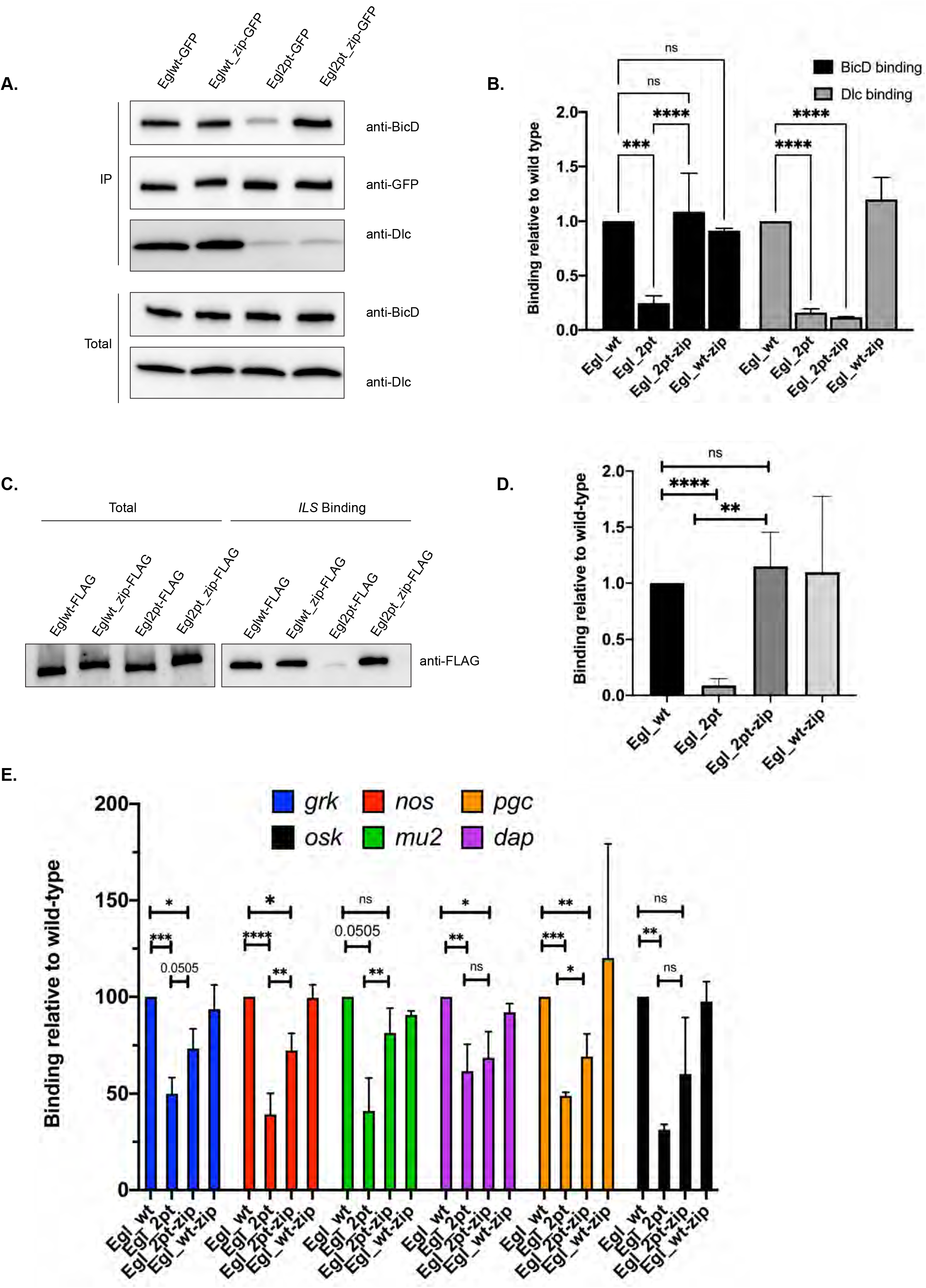
Artificial dimerization of Egl. **(A)** A co-immunoprecipitation experiment was performed using ovarian lysates from flies expressing Egl_wt-GFP, Egl_wt-zip-GFP, Egl_2pt-GFP, and Egl_2pt-zip-GFP. The lysates were incubated with GFP-Trap beads and after binding and wash steps, the bound proteins were eluted in SDS buffer. Eluates were analyzed by western blotting using the indicated antibodies. Total and IP fractions are shown. **(B)** The data from three independent co-immunoprecipitation experiments were quantified. Binding is shown relatively to the BicD and Dlc value obtained with the Egl_wt sample. Neither Egl_2pt nor Egl_2pt-zip is able to associate with Dlc. However, as a consequence of artificial dimerization, the binding to BicD is restored with Egl_2pt-zip. **(C)** Ovarian lysates from the indicated strains were used in an in vitro RNA binding assay. The lysates were incubated with Streptavidin beads coated with the *ILS* localization element. Bound proteins were eluted and analyzed by western blotting. Total and bound fractions are shown. **(D)** Data from three independent *ILS* binding experiments were quantified. Binding was normalized to the level observed in the Egl_wt sample. Using this in vitro assay, artificial dimerization of Egl restores the mRNA binding defect. **(E)** Ovarian lysates from the indicated strains were prepared and the tagged proteins were immunoprecipitated using FLAG antibody beads. Bound mRNAs were extracted and analyzed by reverse transcription followed by quantitative PCR (RTqPCR) using primers against *grk*, *nos*, *mu2*, *dap, pgc* and *osk*. RNA enrichment was normalized relative to the amount of *gamma tubulin* and *rp49* mRNAs that co-precipitated with the beads. Binding was normalized to the level of mRNA enrichment detected with Egl_wt. Using this assay, artificial dimerization of Egl only partially restored the RNA binding deficit. For the BicD/Dlc interaction, a two-way ANOVA with Tukey’s multiple comparison correction was used. For the *ILS* binding experiment, an Unpaired *t-*test was used and for the RTqPCR experiment, a one-way ANOVA with Tukey’s multiple comparison correction was used. ****p<0.0001, *** p<0.001, ** p<0.01, *p<0.05, ns = not significant.

We next examined RNA binding using a well characterized assay (Dienstbier et al., 2009). This assay involves tethering RNA localization elements to beads using a streptavidin-binding aptamer (Dienstbier et al., 2009). Beads bound with the *ILS* localization element from the *I factor* retrotransposon were incubated with ovarian lysates from flies expressing the indicated constructs. Consistent with our published results (Goldman et al., 2019), Egl_2pt displayed reduced RNA binding in comparison to wild-type Egl (Fig. 2C, D). By contrast, using this assay, Egl_2pt-zip bound to the *ILS* localization element at the same level as Egl_wt (Fig. 2C, D). Similar results were obtained using localization elements from *K10* mRNA (*TLS*) and *grk* mRNA (*GLS*) (Supplemental Fig. 1 A, B).

In order to examine the ability of the Egl to associate with native mRNAs, we immunoprecipitated the respective constructs from ovarian lysates, extracted bound mRNAs, and determined their abundance using reverse transcription followed by quantitative PCR (IP RT-qPCR). For this analysis, we determined the level of *gurken* (*grk*), *nanos* (*nos*), *polar granule component* (*pgc*), *mutator2* (*mu2*), *dacapo* (*dap*) and *oskar* (*osk*) that co-precipitated with Egl. The binding of these cargoes were normalized to *gamma tubulin* and *rp49* mRNA, both of which precipitate non-specifically with beads (Supplemental table 1). In comparison to Egl_wt, all six cargoes displayed reduced binding to Egl_2pt (Fig. 2E). In addition, with all six cargoes, partial restoration of binding was observed using Egl_2pt-zip (Fig. 2E). However, the rescue was not complete (Fig. 2E). Thus, using this assay, artificial dimerization of Egl does not completely rescue the mRNA binding function of Egl_2pt.

It was somewhat surprising that the in vitro and in vivo assays produced slightly different results. The most direct comparison can be made using the *GLS* localization element from *grk* mRNA. Using the in vitro aptamer-based assay, Egl_2pt displays a ten fold mRNA binding deficit for this localization element, whereas in the context of the native mRNA, the binding deficit is closer to two fold. In addition, although full rescue of RNA binding is observed with Egl_2pt-zip using the in vitro assay, this is not the case in vivo (Supplemental Fig. 1B and Fig. 2E). As will be shown in subsequent sections, mRNA localization and oocyte specification phenotypes are more consistent with the in vivo (IP RT-qPCR) result.

### Efficient mRNA localization required Egl dimerization

We next examined the localization of *grk* and *mu2* mRNA in mid-stage egg chambers. For this experiment, endogenous Egl was depleted using the afore mentioned early stage driver (maternal alpha tubulin-Gal4; Bloomington stock 7063). This bypasses the function of Egl in the germarium and early stage egg chambers. The same driver was also responsible for expressing the transgenic constructs. *grk* mRNA is localized at the dorsal anterior corner of stage10 egg chambers (Neuman-Silberberg and Schupbach, 1993). This pattern was observed in egg chambers expressing Egl_wt or Egl_wt-zip (Fig. 3A, data not shown). *grk* mRNA was still enriched at the dorsal-anterior corner of egg chambers expressing Egl_2pt. However, the level of enrichment was reduced in comparison to the wild-type control (Fig. 3B, G). This phenotype was only partially rescued in flies expressing Egl_2pt-zip (Fig. 3C, G).

**Figure 3:**
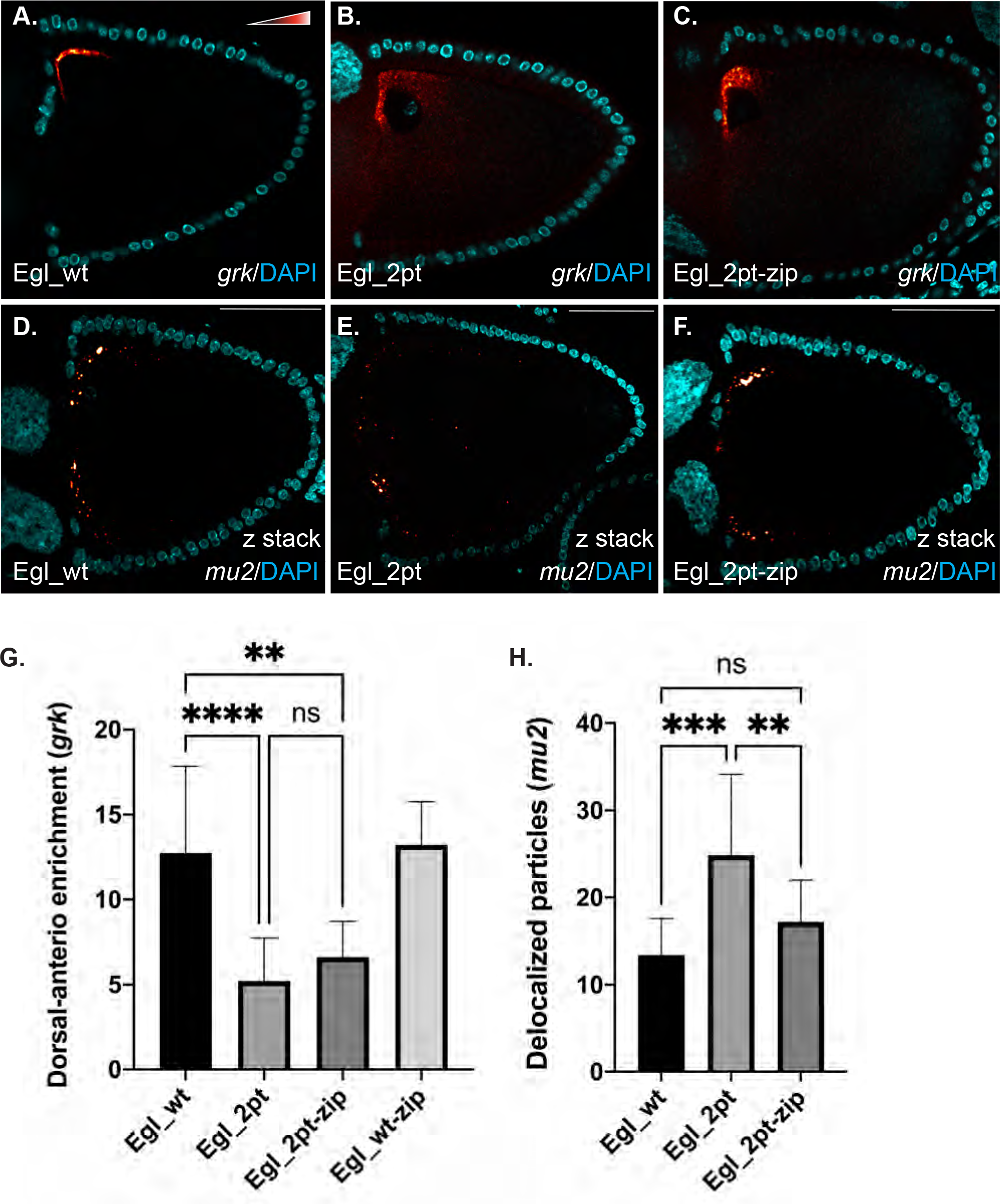
mRNA localization in mid-stage egg chambers. **(A-H)** Ovaries were dissected and fixed from flies expressing the indicated constructs and processed using probes against *grk* (A-C) and *mu2* (D-F). Stage10 egg chambers were imaged for these experiments. The in situ signal is depicted using the Red to White look up table (LUT). The egg chambers were also counterstained with DAPI (cyan). Panels D-F represent a maximum projection of a 8 micron section centered around the mid-saggital plane of the egg chamber. *grk* mRNA localized to the dorsal anterior corner in strains expressing Egl_wt. Although a similar pattern was observed in egg chambers expressing Egl_2pt and Egl_2pt-zip, the level of dorsal-anterior enrichment was reduced (G). *mu2* mRNA localized as distinct particles at the anterior margin of stage10 egg chambers in flies expressing Egl_wt or Egl_2pt-zip. However, in flies expressing Egl_2pt, numerous delocalized particles were observed (H). One-way ANOVA with Tukey’s multiple comparison correction was used. For the data in graph G, ten egg chambers were quantified for each genotype; for the graph in panel H, 10 egg chambers were quantified for Egl_wt and 15 egg chambers were quantified for Egl_2pt and Egl_2pt-zip. ****p<0.0001, *** p<0.001, ** p<0.01, ns = not significant. The scale bar is 50 microns.

*mu2* mRNA has been shown to localize to the anterior cortex of stage10 egg chambers (Kasravi et al., 1999). However, using single molecule fluorescent in situ hybridization (smFISH), we recently demonstrated that *mu2* mRNA localizes to numerous discrete puncta along the anterior margin of the oocyte (Goldman et al., 2021). This pattern was observed in flies expressing Egl_wt or Egl_wt-zip (Fig. 3D, data not shown). In flies expressing Egl_2pt, *mu2* puncta were often delocalized from the anterior margin of the oocyte (Fig. 3E, H). The anterior localization of this mRNA was mostly restored in flies expressing Egl_2pt-zip (Fig. 3F, H). Thus, efficient localization of *grk* and *mu2* mRNA in stage10 egg chambers requires Egl dimerization. Artificial dimerization of Egl_2pt partially (*grk*) or almost completely (*mu2*) restores the localization of these mRNAs.

BicD and Glued (a component of the Dynactin complex) co-localized on microtubules in egg chambers treated with PIPES, indicating active Dynein mediated transport of BicD-linked cargo (Fig. 4A). By contrast, although Glued still localizes to filaments in Egl_2pt egg chambers, BicD was more diffusely localized (Fig. 4B), suggesting defective transport of BicD-linked cargo. Filament localization was only partially rescued in egg chambers expressing Egl_2pt-zip (Fig. 4C). Thus, although artificial dimerization is able to restore the Egl-BicD interaction, this complex does not efficiently associate with Dynein. mRNAs such as *grk* may be more reliant on Dynein mediated transport for its correct localization in comparison to mRNAs such as *mu2*. This might explain why artificial dimerization of Egl_2pt only partially restored *grk* localization whereas *mu2* localization was more fully rescued.

**Figure 4:**
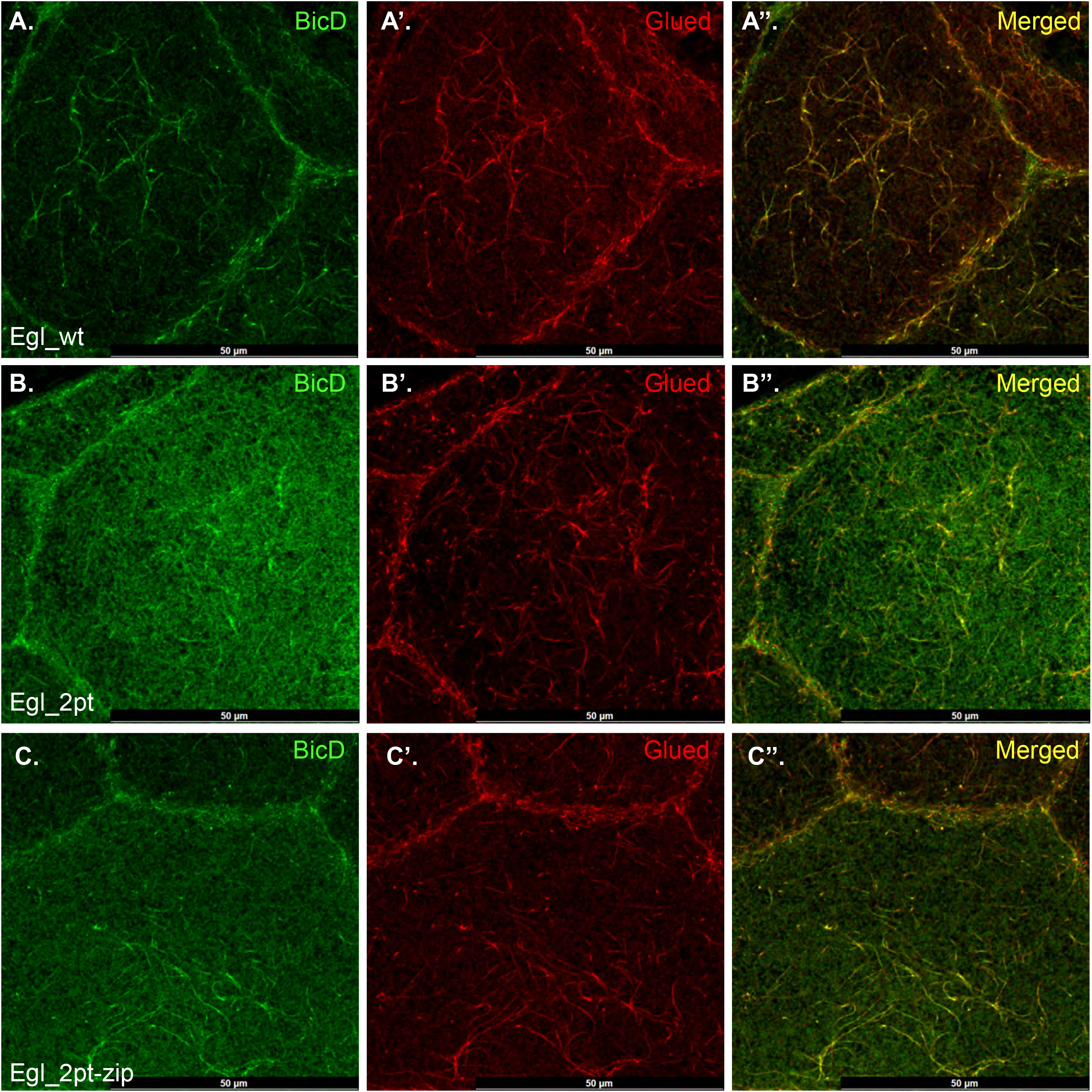
Localization of BicD to microtubule filaments. **(A-C)** Egg chambers from the indicated genotypes were incubated with PIPES prior to fixation. Egg chambers were processed for immunofluorescence using antibodies against BicD (green) and Glued (red). A merged image is also shown. BicD and Glued co-localize on microtubule filaments in strains expressing Egl_wt (A). By contrast, BicD did not display a filamentous localization pattern in Egl_2pt egg chambers (B). Egl_2pt-zip produced an intermediate phenotype and partial filament localization of BicD could be observed (C). The scale bar is 50 microns.

### Egl dimerization is required for meiosis restriction and oocyte specification

In addition to its role in mRNA localization in mid-stage egg chambers, Egl has additional functions in the germarium. Null alleles of *egl* display meiotic defects and fail to specify an oocyte (Carpenter, 1994). A similar phenotype is observed in *egl_2pt* mutants and in *egl* mutants that are specifically defective in mRNA binding (Goldman et al., 2021; Navarro et al., 2004). We therefore wished to test whether artificial dimerization of Egl_2pt could rescue these functions of Egl in the germarium and early stage egg chambers.

Oocyte specification occurs within the germarium. A cystoblast is produced at the distal tip of the germarium via division of a germline stem cell. The cystoblast replicates four times to generate a cyst containing sixteen interconnected cells. Within this early cyst, two cells initiate a meiotic program as evident by the formation of the synaptonemal complex. As the cyst matures, one of these cells will be specified as the oocyte and will retain the synaptonemal complex. The other cell will eventually exit meiosis and revert to a nurse cell fate (Hughes et al., 2018; Huynh and St Johnston, 2004) (Fig. 5A). In *egl* nulls, numerous cells within the cyst initiate a meiotic program and form a synaptonemal complex. However, this effect is temporary; meiosis is eventually terminated in all cells of the cyst and an oocyte is never specified (Carpenter, 1994; Huynh and St Johnston, 2000).

**Figure 5:**
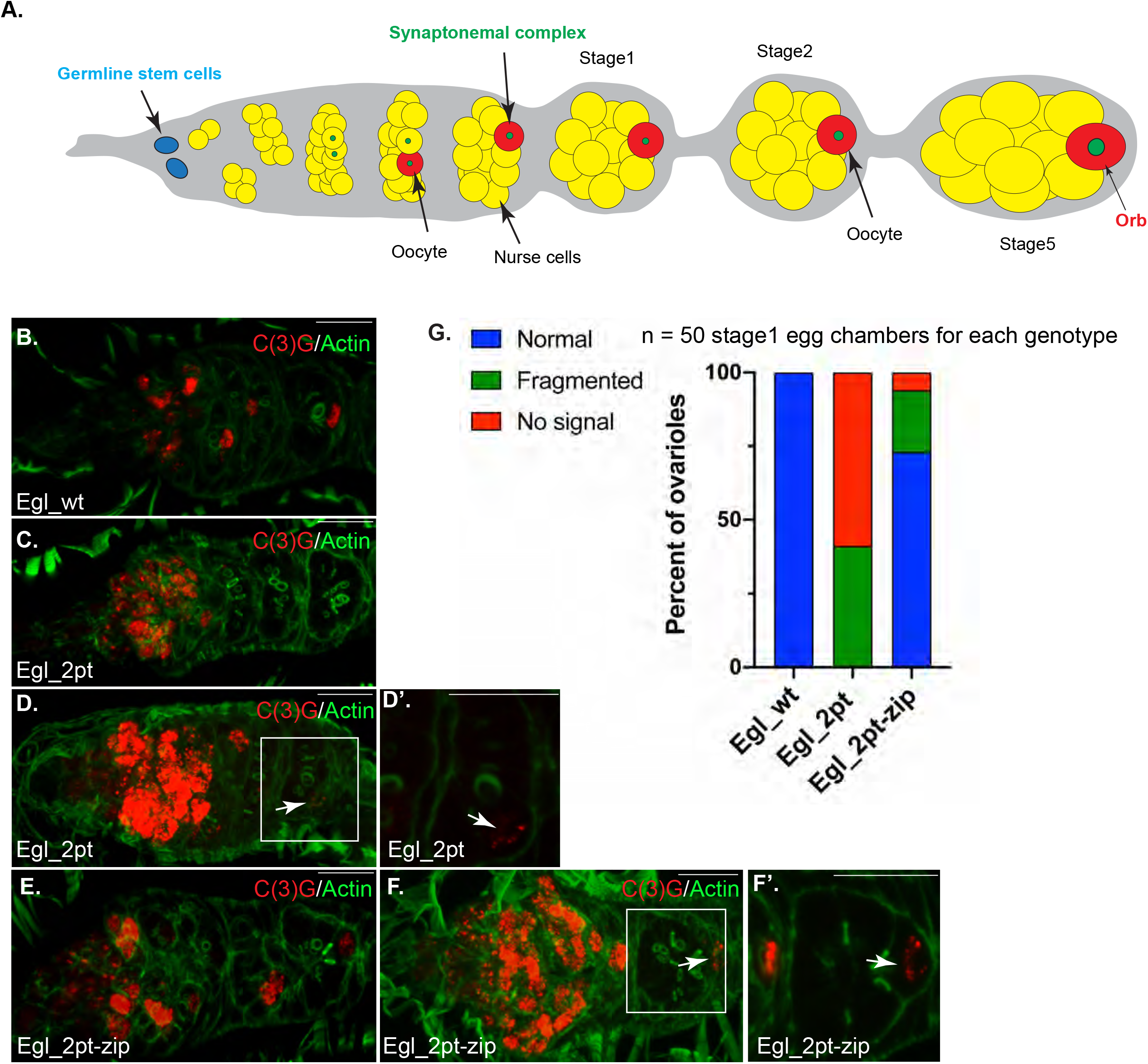
Artificial dimerization partially restores the meiosis defect of Egl_2pt mutants. **(A)** A schematic of the *Drosophila* germarium and early stage egg chambers. Germline stem cells, the synaptonemal complex, and the localization of Orb are indicated. **(B-G)**. Ovaries from flies expressing Egl_wt, Egl_2pt, or Egl_2pt-zip were fixed and processed for immunofluorescence using an antibody against C(3)G, a marker for the synaptonemal complex (red). The tissues were also counter-stained with Phalloidin to reveal the actin cytoskeleton (green). In flies expressing Egl_wt, 100% of ovarioles contained C(3)G staining within one cell of stage1 egg chambers (B, G). By contrast, ovarioles from Egl_2pt mutants either lacked C3G staining in stage1 egg chambers and beyond (C, G), or displayed residual puncta of C(3)G (D, G). D’ is a magnified view of the image in D. In flies expressing Egl_2pt-zip, the vast majority displayed normal C(3)G staining in stage1 egg chambers (E, G). However, in 21% of ovarioles, the C(3)G staining in stage1 egg chambers was fragmented (F, G). F’ is a magnified view of the image in F. The scale bar is 10 microns.

In order to examine this process in our mutants, germaria were stained using an antibody against C(3)G, a component of the synaptonemal complex (Page and Hawley, 2001). Endogenous Egl was depleted in the germanium using the *nanos*-Gal4 driver (Bloomington stock #4937). The same driver was also responsible for expressing transgenic wild-type or mutant constructs. A phenotype similar to the null was observed in *egl_2pt* mutants. Like the null allele, in *egl_2pt* mutants, numerous cells within the cyst inappropriately entered meiosis. However, as with the nulls, this effect was transient, and by stage1, the synaptonemal complex was either undetected or present as faint residual puncta (Fig. 5B, C, D, D’, G). This phenotype was significantly restored in flies expressing Egl_2pt-zip and 73% of germaria displayed normal synaptonemal complex formation in stage1 egg chamber (Fig. 5E, G). However, the rescue was not complete and 21% of egg chambers displayed fragmented C(3)G staining (Fig. 5F, F’, G). Based on these results, we conclude that proper progression of meiosis requires Dlc-mediate Egl dimerization and that artificial dimerization Egl is able to partially compensate for loss of Dlc binding.

We next examined the specification of oocyte fate by determining the localization of Orb in the germarium and early stage egg chambers. Orb, the *Drosophila* ortholog of Cytoplasmic Polyadenylation Element Binding Protein 1, is enriched within the two cells that initially enter meiosis. Subsequently, Orb becomes highly enriched within the oocyte, and is therefore a useful marker of oocyte fate specification and maintenance (Huynh and St Johnston, 2000; Lantz et al., 1994). In contrast to egg chambers expressing Egl_wt or Egl_wt-zip, Orb is never localized to a single cell with the germaria of Egl_2pt mutants (Fig. 6A, B, D, G). This phenotype is partially restored in Egl_2pt-zip flies, and about 30% of ovarioles proceed though normal development (Fig. 6C, F, G). However, in about 60% of ovarioles, the oocyte is either not specified, or if specification does occur, oocyte fate is not maintained (Fig. 6E, G). Thus, Dlc-mediated dimerization of Egl is required for specification and maintenance of oocyte fate, and artificial dimerization of Egl is only able to partially rescue this phenotype.

**Figure 6:**
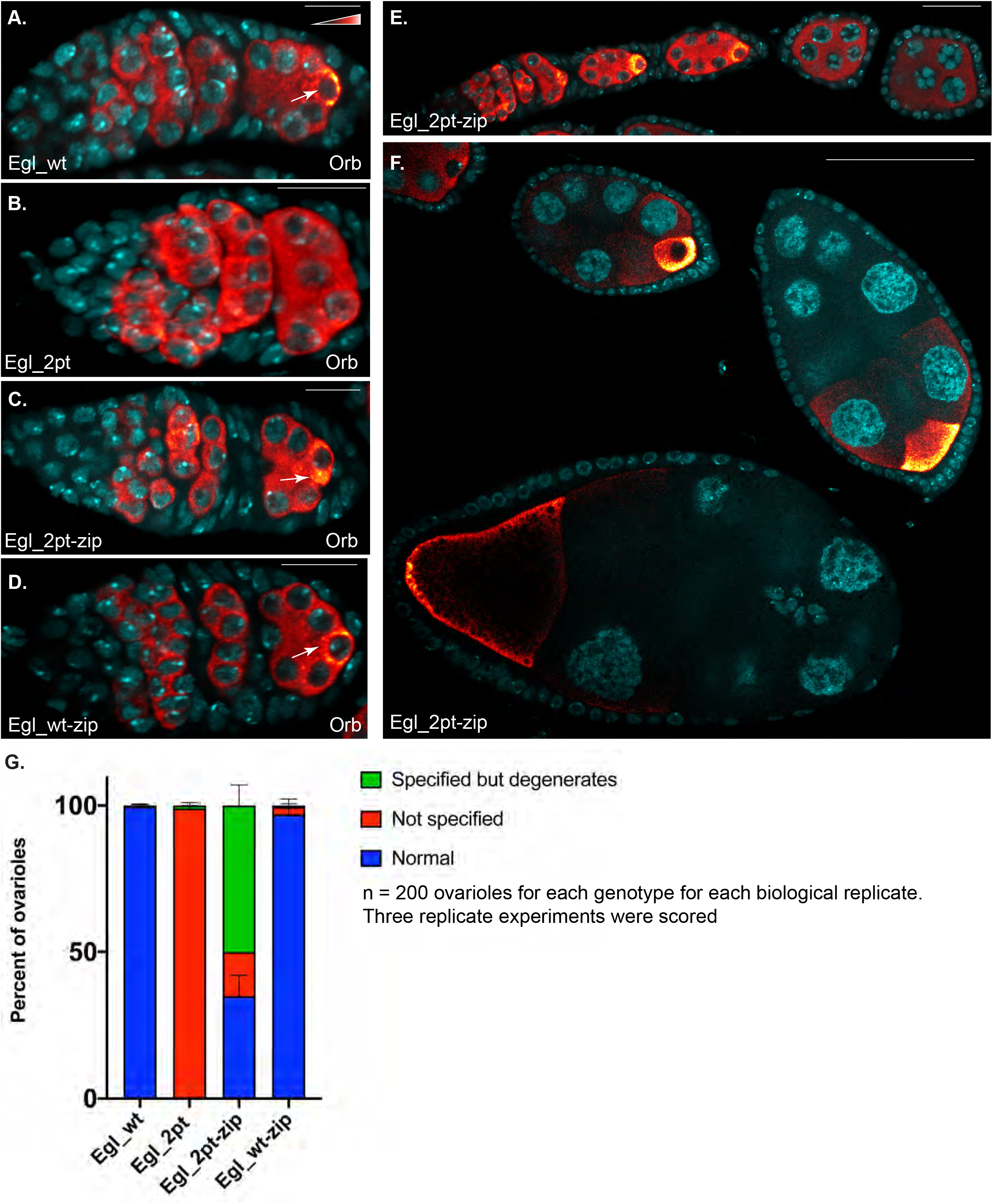
Oocyte specification and maintenance. **(A-G)** Ovaries were dissected and fixed from flies expressing Egl_wt, Egl_2pt, Egl_2pt-zip or Egl_wt-zip. The ovaries were processed for immunofluorescence using an antibody against Orb, a protein that is enriched within the oocyte of stage1 egg chambers and beyond (indicated by arrows in A, C and D). The signal for Orb is shown using a Red to White look up table (LUT). Low intensity pixels are depicted in red whereas high intensity pixels are shown in white. In contrast to egg chambers expressing Egl_wt and Egl_wt-zip, Orb was never localized to a single cell in egg chambers expressing Egl_2pt (A, B, D, G). In flies expressing Egl_2pt-zip, three phenotypes were observed (G). In a small percentage of ovarioles, Orb was not localized to a single cell within stage1 egg chambers. In the remainder of the ovarioles, Orb was either correctly localized throughout egg chamber maturation (F, G) or was initially localized correctly and then became delocalized once oocyte fate was lost (E, G). Thus, artificial dimerization of Egl_2pt partially rescues the oocyte specification and and maintenance defect. The scale bar in A-D is 10 microns. In panel E the scale bar is 20 microns, and 50 microns in panel F.

### Early-stage mRNA localization is highly dependent on Egl dimerization

Several mRNA’s that are localized in an Egl-dependent mechanism are known to be enriched within the presumptive oocyte (Kasravi et al., 1999; Roth et al., 1995). For this analysis, we examined the localization of *grk* and *mu2* within the germarium and early stage egg chambers. Both cargoes were highly enriched within a single cell of stage1 and stage2 egg chambers in files expressing Egl_wt (Fig. 7A, A’, D, D’ arrow). A similar pattern was observed in Egl_wt-zip egg chambers (Supplemental Fig. 1E, F). By contrast, both mRNAs were completely delocalized in Egl_2pt mutants (Fig. 7B, B’, E, E’). In flies expressing Egl_2pt-zip, a modest enrichment of *grk* and *mu2* mRNA could be detected within the oocyte of stage1 and stage2 egg chambers (Fig. 7C, C’, F, F’, dashed outline). However, quantification of oocyte enrichment revealed that this rescue was only partial (Fig. 7G, H). Thus, oocyte localization of these mRNAs requires Dlc-mediated Egl dimerization. Artificial dimerization is only able to partially substitute for Dlc.

**Figure 7:**
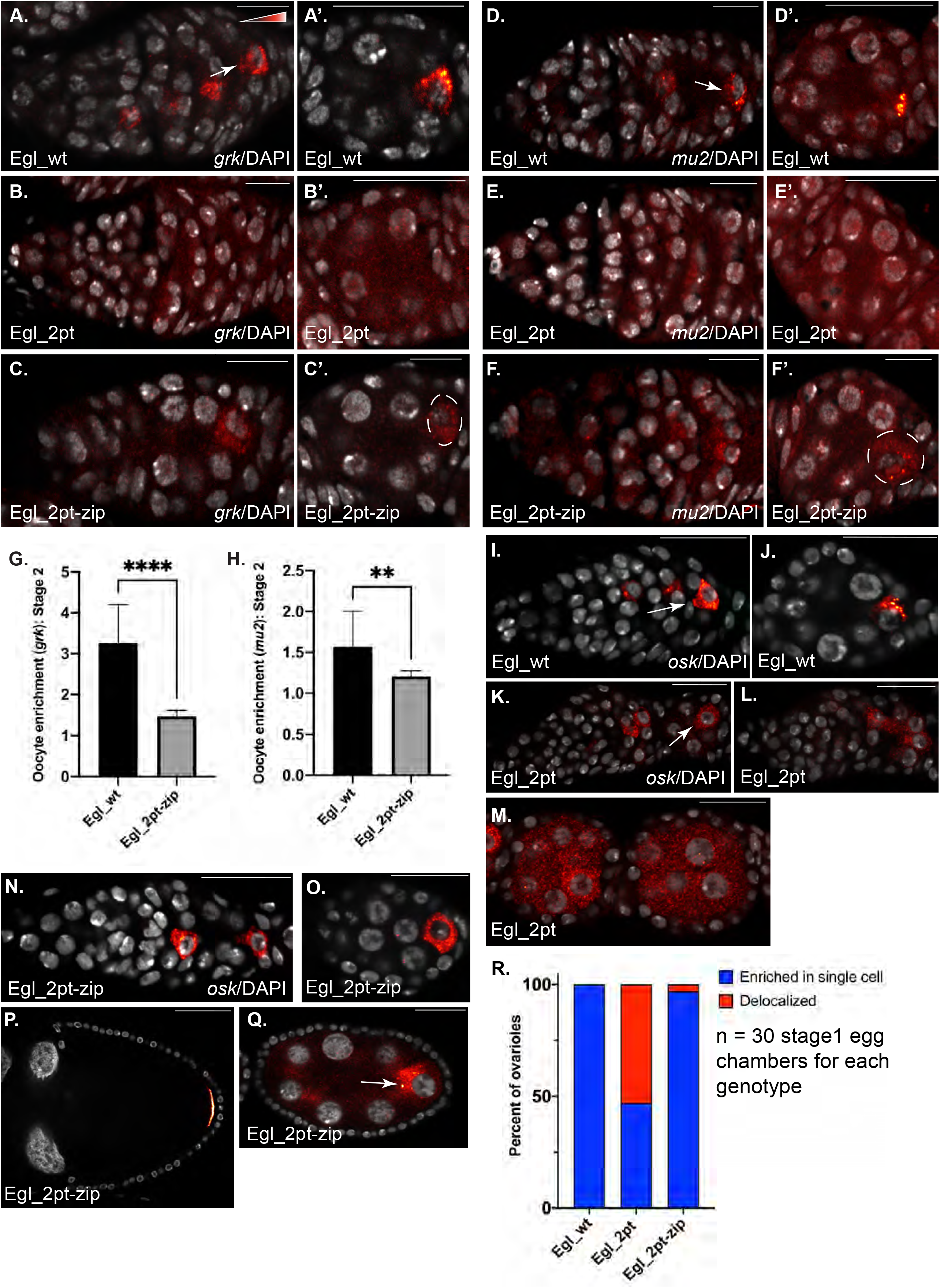
mRNA localization in early-stage egg chambers. **(A-H)** Ovaries were dissected and fixed from flies expressing the indicated constructs. The germaria and early stage egg chambers were processed for single molecule fluorescent in situ hybridization (smFISH) using probes against *grk* (A-C) or *mu2* (D-F). The egg chambers were also counterstained with DAPI (shown in greyscale). The in situ signal for *grk* and *mu2* is shown using the same Red to White look up table (LUT) that was used for Orb. The oocyte in C’ and F’ is indicated with a dashed line. *grk* and *mu2* mRNA are highly enriched in the oocyte in stage1 and 2 wild-type egg chambers but are completely delocalized in Egl_2pt mutants. Both mRNA are partially enriched with the oocyte of flies expressing Egl_2pt-zip but are reduced in comparison to the wild-type (G, H). An unpaired *t*-test for used for the analysis show in G and H. ****p<0.0001, *** p<0.001. Twelve egg chambers were quantified for each genotype depicted in graphs G and H. **(I-R)** The same strains were processed using smFISH probes against *osk*. *osk* mRNA is enriched within a single cell in wild-type stage1 and 2 egg chambers (I, J, R). In 47% of stage1 egg chambers expressing Egl_2pt, *osk* could be detected within a single cells (K, R). In the remainder, *osk* mRNA was delocalized (L, R). *osk* was delocalized in stage3 and later egg chambers in flies expressing Egl_2pt (M). In flies expressing Egl_2pt-zip, osk was normally localized in egg chambers that retained oocyte fate (N, O, P, R). However, even in egg chambers that had lost oocyte fate, osk enrichment could often be detected within a single cell (Q). The scale bar in A-F is 10 microns; in I-Q, the scale bar is 20 microns and in panel P, the scale bar is 50 microns.

We next examined the localization of *osk* mRNA within the germarium and early stage egg chambers. Although *osk* is localized at the posterior pole in mid-stage egg chambers by the plus-end directed microtubule motor, Kinesin-1 (Brendza et al., 2000), its transport into the oocyte is though to be dependent on Dynein (Jambor et al., 2014; Ryu et al., 2017). As expected, *osk* mRNA was localized to a single cell in stage1 and stage2 egg chambers in flies expressing Egl_wt (Fig. 7I, J, R). Unexpectedly, and in contrast to *grk* and *mu2*, *osk* mRNA was restricted to a single cell in 47% of stage1 egg chambers from flies expressing Egl_2pt (Fig. 7K, L, R). This is noteworthy because neither meiosis nor oocyte markers such as Orb are enriched within a single cell in stage1 egg chambers in these mutants (Figs. 5, 6). The accumulation of *osk* mRNA within this single cell might indicate that this cell still retains some characteristics of an oocyte. However, by stage3, *osk* is no longer localized to a single cells in flies expressing Egl_2pt (Fig. 7M). In flies expressing Egl_2pt-zip, *osk* localizes normally in egg chambers that specify and maintain oocyte fate (Fig. 7N-P). In addition, even in egg chambers that have lost oocyte fate, *osk* is occasionally enriched within a single cell (Fig. 7Q, arrow). This cell presumably corresponds to the dedifferentiating oocyte. Thus, among these mRNA cargoes, the localization of *osk* to a single cell is the least reliant on Egl dimerization.

Similar to the *osk* mRNA phenotype, Dynein was less dependent on Egl’s ability to dimerize for its oocyte enrichment. Dynein could be detected within a single cells in stage1 egg chambers in flies expressing Egl_2pt (Supplemental Fig. 1C). However, this enrichment was reduced in stage2 egg chambers, coincident with loss of oocyte fate (Supplemental Fig. 1C’). In egg chambers expressing Egl_2pt-zip, Dynein was either normally localized within the oocyte (Supplemental Fig. 1D, D’, arrow) or became delocalized in those egg chambers in which oocyte fate was lost (Supplemental Fig. 1D, asterisk). Thus, it appears that as long as oocyte fate is maintained, Dynein is able to localize normally within the oocyte. A similar phenotype was observed for BicD. In Egl_2pt-zip egg chambers in which oocyte specification occurred normally, BicD and Egl were co-localized within the oocyte (Supplemental Fig.1 G, H). However, if oocyte fate was lost, neither BicD nor Egl were enriched within a single cell (Supplemental Fig. 1I).

In conclusion, Dlc-dependent dimerization of Egl is required for several processes during oogenesis. Within the germarium, Egl dimerization is required for meiosis to proceed normally, for specification of the oocyte, and for localization of mRNAs such as *grk* and *mu2* to the oocyte. Artificial dimerization of Egl is not capable of fully rescuing these early phenotypes. In mid-stage egg chamber, mRNA localization is less affected by Egl dimerization. Even in the absence of Egl-dimerization, *grk* and *mu2* mRNA remained relatively localized. Distinct mechanisms of localization might be active in early and mid stage egg chambers. If true, this would account for the different localization phenotypes observed for *grk* and *mu2* mRNA in early vs mid-stage egg chambers. Future studies will be critical to reveal these potentially distinct mechanisms.

## DISCUSSION

Navarro and colleagues demonstrated several years ago that Egl contains a Dlc-interaction motif within its C-terminus (Navarro et al., 2004). They further showed that point mutations within this region, which they referred to as *egl_2pt*, resulted in oocytes specification defects (Navarro et al., 2004). At that time, the role of Dlc in this pathway was unclear, and it was suggested that Dlc might serve as a way to link Egl and its associated mRNA cargo with the Dynein motor. Subsequent research has shown that the actual link between Egl and Dynein is the adaptor protein, BicD (McClintock et al., 2018; Sladewski et al., 2018). We recently demonstrated that the function of Dlc in this pathway is to facilitate dimerization of Egl, a pre-requisite for efficient mRNA binding (Goldman et al., 2019).

### The role of Egl dimerization

Although initially identified as a component of the Dynein motor, Dlc (also known as LC8) has been shown to interact with numerous proteins. In fact, three hundred nineteen interactions are listed for human LC8 and sixty three interactions are listed for *Drosophila* Dlc on BioGRID. The emerging picture suggests that the main role for Dlc is to facilitate the dimerization of proteins, in particular those that harbor regions of low complexity or intrinsically disordered stretches (Barbar, 2008; Reardon et al., 2020). Apart from its central RNA binding domain, Egl contains large stretches of low complexity regions and a single consensus Dlc-interaction motif at its C-terminal end (Navarro et al., 2004). Although structural information is not yet available regarding the mechanism of Egl dimerization, it is tempting to speculate that Dlc-mediated dimerization alters the conformation of Egl’s RNA binding domain in such a way as to promote efficiently cargo binding.

Egl is thought to be the primary mRNA adaptor for the Dynein motor in *Drosophila* oocytes and embryos (Dienstbier et al., 2009; Vazquez-Pianzola et al., 2017). As such, it has to be able to bind diverse cargo. In select cases, it has been demonstrated that Egl recognized a secondary structure in its target mRNAs (Dienstbier et al., 2009; Vazquez-Pianzola et al., 2017). However, whether comparable structures are present in all Egl mRNA cargoes is not known. Thus, having an RNA binding domain that is somewhat flexible and unstructured might be required for Egl to recognize diverse cargo. Dlc-mediated dimerization might in turn promote a particular structural conformation such that only mRNAs destined for localization are coupled to Dynein.

Our results indicate that artificial dimerization of Egl is able to partially substitute for Dlc in this pathway. The reason for this incomplete rescue might have to do with the three dimensional conformation of the complex. In our construct, the leucine zipper is placed at the C-terminus of Egl, and therefore dimerization is induced at this site. The structure of the resulting complex might not be identical to the dimeric structure that is produced via Dlc-mediated dimerization. As such, the RNA binding domain of Egl might not adopt an optimal conformation, thus resulting in inefficient cargo binding and Dynein mediated transport.

Another potential reason for incomplete rescue might have to do with the leucine zipper itself. The zipper used in these studies is from the yeast transcription factor, GCN4. In the yeast protein, this zipper functions a homodimerization domain (Abel and Maniatis, 1989; Hope and Struhl, 1987) and we assume that the same properties apply in our system. However, the possibility exists that Egl_2pt-zip might instead dimerize with *Drosophila* proteins that also contain a leucine zipper. There are twenty four proteins encoded in the fly genome that are predicted to contain a leucine zipper. We have shown previously, that Egl_2pt-zip restores the homodimerization of Egl (Goldman et al., 2019). However, if Egl_2pt-zip also heterodimerizes with one of these other leucine zipper-containing proteins, this would titrate away Egl_2pt-zip from functional transport complexes, resulting in incomplete rescue.

Recently, Panoramix (Panx), a protein that functions in the piRNA pathway was also shown to dimerize in a Dlc-dependent manner (Eastwood et al., 2021; Schnabl et al., 2021). Artificial dimerization of a *panx* mutant that could no longer interact with Dlc (using the same GCN4 leucine zipper) restored the function of *panx* (Eastwood et al., 2021; Schnabl et al., 2021). Similar to our findings, the rescue of certain *panx* mutant phenotypes was incomplete (Schnabl et al., 2021).

### RNA binding by Egl

The RNA binding activity of Egl is usually examined using either the in vitro aptamer based assay or in vivo using IP RT-qPCR. We were surprised to find that these assays reported different results regarding RNA binding activity when mutant versions of Egl were used. For instance, using the in vitro assay, Egl_2pt has negligible RNA binding activity, yet using IP RT-qPCR, the RNA binding deficit is only about two-fold (Fig. 2C, E). The mid-stage mRNA localization defects observed in this mutant background (Fig. 3) are more in line with the two-fold RNA binding deficit rather than the near complete loss of RNA binding.

We observed a similar phenotype using Egl mutants containing point mutations in the RNA binding domain (Goldman et al., 2021). The Egl_rbd3 mutant contains eight amino acid substitutions, whereas Egl_rbd6 contains just three of these eight substitutions. Using the in vitro assay, both mutants had minimal RNA binding activity. However, Egl_rbd3 was on average ten-fold defective for binding to native mRNAs in vivo whereas Egl_rbd6 was only about two-fold defective (Goldman et al., 2021). One possible reason for this difference might have to do with the highly structured part of the RNA that corresponds to the Streptavidin aptamer (Srisawat and Engelke, 2001). The aptamer, by virtue of its structure, might alter the conformation of the *ILS* or other localization sequences in such a way that even a subtle defect in Egl’s RNA binding activity results in dramatic loss of binding. Another possible scenario is that in vivo Egl recognizes not just the RNA localization sequence alone, but also proteins that are associated with the mRNA. These factors may promote the binding of Egl to localizing mRNAs. In this scenario, any inherent defect in Egl’s RNA binding activity may be attenuated by these additional interactions.

Although it is clear that Egl is the primary mRNA adaptor for the Dynein motor in oocytes and embryos, there is still much we do not know regarding the mechanism by which Egl is able to recognize its diverse cargoes. For instance, Egl_rbd6 is compromised for binding to cargoes such as *grk*, *nos*, and *mu2*. However, this mutant still retained binding activity for cargoes such as *bcd* and *hts* (Goldman et al., 2021). Thus, not all cargoes are equivalent when it comes to binding Egl. Additional studies will be required to fully understand the mechanism of Egl-mRNA binding.

### Transport of *osk* mRNA into the oocyte

The complex responsible for transporting *osk* mRNA to the posterior pole of the oocyte involves an adaptor protein and the plus-end directed microtubule motor, Kinesin-1 (Brendza et al., 2000; Gaspar et al., 2017; Veeranan-Karmegam et al., 2016). However, transport of *osk* from the nurse cells into the oocyte relies on Dynein. Evidence for this comes from the finding that specific sequences within the 3’UTR of *osk* are required for nurse cell to oocyte transport (Jambor et al., 2014; Ryu et al., 2017). In addition, *osk* mRNA can be detected in complex with the Egl/Dynein motor (Fig. 2E) (Sanghavi et al., 2016). However, the Macdonald lab recently demonstrated that Egalitarian competes with another RNA binding protein, Staufen, for binding to this region of *osk* mRNA. The competition between these pathways is necessary for *osk* to be transported into the oocyte, but also weaken its interaction with the Dynein complex, such that once in the oocyte, plus-end directed microtubule transport can occur (Mohr et al., 2021). In contrast to *osk* transport in mid and late-stage egg chambers, not much is known regarding the mechanism by which *osk* mRNA is localized in the germarium and early stage egg chambers. Consistent with published findings (Little et al., 2015), we observed that *osk* mRNA was restricted to a single cell in stage1 egg chambers (Fig. 7I). However, unlike cargoes such as *mu2* and *grk* that are transported by Dynein for the entire course of their localization pathway, localization of *osk* mRNA was less affected by loss of Egl dimerization (Fig. 7K, R). Surprisingly, even once egg chambers had lost oocyte fate, an accumulation of *osk* could often still be detected within a single cell (Fig. 7Q, arrow). At present, we do not have a definitive explanation for this phenotype. One possibility is that a different motor is responsible for the early-stage transport of *osk* mRNA into the oocyte. It is also possible that *osk* associates with the Dynein motor in an Egl-dependent and Egl-independent manner. A final possibility, and one that we favor, is that *osk* transport into the oocyte relies on Egl and Dynein. However, once in the oocyte, *osk* mRNA is more stable than mRNAs such as *grk* and *mu2*. Thus, even once the oocyte dedifferentiates in these mutants, a residual accumulation of *osk* can still be detected within this cell.

## ACKNOWLEDGEMENTS

We are grateful to Simon Bullock for providing the *ILS* and *TLS* aptamer constructs and Scott Hawley for providing the C(3)G antibody. We would also like to thank Vladimir Gelfand for providing the Glued antibody. We are grateful to the Bloomington Stock Center, Developmental Studies Hybridoma Bank, and the *Drosophila* Genomics Resource Center for providing fly strains, antibodies, cell lines, and DNA constructs. This work was supported by a grant from the National Institutes of Health to G.B.G (R01GM100088).

## MATERIALS AND METHODS

### Fly stocks and DNA constructs

The following shRNA stocks were used:

*egl* shRNA-1 (Bloomington stock center; #43550, donor TRiP).

shRNA expression was driven using:

P{w[+mC]=matalpha4-GAL-VP16}V37 (Bloomington Stock Center, #7063; donor Andrea Brand) for early-stage expression.

P{w[+mC]=GAL4::VP16-nos.UTR}CG6325[MVD1] (Bloomington Stock center, #4937; donor Ruth Lehmann) for expression in the germanium.

Vectors expressing Egl_wt and Egl_2pt were designed in a previous study (Goldman et al., 2019). The expression of these constructs were driven using a maternal promoter. For this study, the coding regions from those constructs were sub-cloned into the the pUASp-attB-K10 vector (Koch et al., 2009) containing either a C-terminal GFP tag or 3xFLAG tag. The leucine zipper from yeast GCN4 (AAL09032.1) along with a preceding flexible linker sequence was synthesized with *Drosophila* codon optimization by Genewiz. The zipper was cloned downstream of the GFP or FLAG tag. The GFP tagged transgenic constructs were inserted at the ZH-86Fa site (Bloomington stock center; #24486, donor Konrad Basler & Johannes Bischof) and the FLAG tagged constructs were inserted at su(Hw)attP1 site (Bloomington stock center; #34760, donor Norbert Perrimon). The strains were injected by BestGene Inc. Fly stocks and crosses used for these experiments were maintained at 250C. The *ILS* and *TLS* aptamers were a gift from Simon Bullock. The *GLS* aptamer construct was created using Gibson assembly (NEB). The *GLS* sequence was synthesized by Genewiz and this double stranded DNA was used to replace the *ILS* sequence in the *ILS* aptamer vector.

### Antibodies

The following antibodies were used: anti-FLAG (Sigma Aldrich, F1804, 1:5000 for western, 1:500 for immunofluorescence), mouse anti-BicD (Clones 1B11 and 4C2, Developmental Studies Hybridoma Bank, 1:30 for immunofluorescence; 1:300 for western, donor R. Steward), rabbit anti-Ctp (Abcam, ab51603, 1:5000 for western), GFP nanobody booster (Chromotek; 1:500 for immunofluorescence), mouse anti-GFP (Clontech, JL-8, 1:5000 for western), rabbit anti-GFP (Chromotek; 1:3000 for immunofluorescence), C3G (generous gift from S. Hawley, mouse anti-C(3)G, 1:500), Orb (mouse anti-Orb, clone 4H8, 1:30 dilution), mouse anti-Dhc (Developmental Studies Hybridoma Bank, clone 2C11-2, 1:50; for immunofluorescence, donor J. Scholey), rabbit anti-Glued (generous gift from V. Gelfand, 1:500 for immunofluorescence). The following secondary antibodies were used: goat anti-rabbit Alexa 594, 555 and 488 (Life Technologies, 1:400, 1:400 and 1:200 respectively); goat anti-mouse Alexa 594, 555 and 488 (Life Technologies, 1:400, 1:400 and 1: 200 respectively) goat anti-mouse HRP (Pierce, 1:5000); and goat anti-rabbit HRP (Pierce, 1:5000).

### RNA binding studies

The in vitro RNA binding experiments using *ILS*, *TLS* and *GLS* localization elements were performed as previously described (Dienstbier et al., 2009; Goldman et al., 2019). In brief, 5μg of RNA was refolded in 10μL of *Drosophila* extraction buffer (DXB: 25mM HEPES pH 6.5, 50mM KCl, 1mM MgCl2, 250mM sucrose, 0.05% NP40, supplemented with 10μM MgATP and 1mM DTT at the time of use). The refolded RNA was next incubated with High Capacity Streptavidin Agarose Beads (Pierce) in 90μL DXB for 1.5 hours at 4°C while nutating. Extracts from frozen *Drosophila* ovaries were prepared using lysis buffer (50mM Tris pH 7.5, 200mM NaCl, 0.2mM EDTA, 0.05% NP40, Halt protease inhibitor cocktail, Pierce). 600ug of total protein was used in each binding experiment. Extracts were incubated with the RNA-bound streptavidin beads for 15 min at room temperature followed by 30 min at 4°C with nutation. The beads were then washed 4 times in lysis buffer, the bound proteins were eluted by boiling in Laemmli buffer and examined by western blotting.

The binding of Egl constructs to native mRNAs from ovaries was analyzed as previously described (Goldman et al., 2019). Briefly, 600ug of lysate was used in each immunoprecipitation. The tagged proteins were immunoprecipitated using FLAG M2 antibody beads (Sigma). Co-precipitating RNAs were reverse transcribed using random hexamers and Superscript III (Life Technologies). Quantitative (qPCR) was performed using SsoAdvanced Universal SYBR Green Supermix (Bio-Rad). A Bio-Rad CFX96 Real-Time PCR System was used for this experiment. Fold enrichment was calculated by comparing ct values for each RNA to that obtained for *γ-tubulin* and *rp49*.

### Protein-protein interaction

A co-immunoprecipitation experiment was used to analyze protein-protein interaction (Goldman et al., 2019). In brief, ovaries were homogenized into lysis buffer (50mM Tris pH 7.5, 200mM NaCl, 0.2mM EDTA, 0.05% NP40 and Halt protease inhibitor cocktail, Pierce) and cleared by centrifugation at 10,000g at 40C for 5min. 1mg of total protein was used per immunoprecipitation. Immunoprecipitation was performed by incubating lysates at 40C for 1hour with GFP-trap beads (Chromotek). Next, the beads were washed four times with lysis buffer. Co-precipitating proteins were eluted in Laemmli buffer, run on a gel, and analyzed by western blotting.

### Immunofluorescence and in situ hybridization

Immunofluorescence and in situ hybridization were performed as previously described (Goldman et al., 2019). Ovaries were dissected and fixed in 4% formaldehyde (Pierce) for 20 min at room temperature. For immunofluorescence experiments, primary antibody was incubated in blocking buffer (PBS + 0.1% Triton X-100 + 2% BSA) overnight at 4°C. The next day, the samples were washed three times in PBST (PBS + 0.1% Triton X-100) and incubated overnight with the fluorescent secondary antibody in the same blocking buffer. The samples were then washed four times with PBST, stained with DAPI, and mounted onto slides using Prolong Diamond (Life technologies). For in situ hybridization, the ovaries were dissected and fixed as above. After fixation, ovaries were stored in in 100% methanol at −20°C for 1 hour. Next, the samples were re-hydrated with three 10 min washes using a solution of PBST and 100% methanol (3:7, 1:1, 7:3) and rinsed four times with PBST. The hydrated samples were next washed for 10 minutes in Wash Buffer (4xSSC, 35% deionized formamide, 0.1% Tween-20). Fluorescent oligo probes (Stellaris probes) were obtained from Biosearch technologies. Fluorescent probes diluted in Hybridization Buffer (10% dextran sulfate, 0.1mg/ml salmon sperm ssDNA, 100 μl vanadyl ribonucleoside (NEB biolabs), 20ug/ml RNAse-free BSA, 4x SSC, 0.1% Tween-20, 35% deionized formamide) were added to the ovaries and incubated overnight at 37°C. The following day, the samples were washed twice with pre-warmed Wash Buffer for 30 min. After two rinses with PBST and two rinses with PBS, the samples were counter-stained with DAPI, and mounted onto slides using Prolong Diamond.

### Microscopy

Images were captured on either a Zeiss LSM 780 inverted confocal microscope or an inverted Leica Stellaris confocal microscope. Images were processed for presentation using Fiji, Adobe Photoshop, and Adobe Illustrator. All imaging experiments were performed at the Augusta University Cell Imaging Core.

### Quantification

All western blot images were acquired on a Bio Rad ChemiDoc MP. Band intensities were quantified using the Bio Rad Image lab software. RNA and protein enrichment was quantified by measuring the average pixel intensity of the localized signal and dividing by the average pixel intensity of the delocalized signal. For quantifying stage2 oocyte enrichment, the localized signal in the oocyte was compared to the average signal in the rest of the egg chamber. In mid-stage egg chamber, the localized signal for *grk* mRNA at the dorsal-anterior corner was compared to the delocalized signal in the rest of the oocyte. In order to stay consistent between images, the focal plan containing the karyosome of the oocyte nucleus was used for *grk* signal quantification. The above quantifications were performed using the Zeiss Zen Black software. For quantification of *mu2* signal, an 8 micron z stack was analyzed for each sample. The mid-saggital plane of the egg chamber was used as the center and 4 focal planes above and below were imaged. A maximum projection image was then generated and delocalized particles were counted. Localized particles were considered to be within 20 microns from the anterior margin of the oocyte. The remainder of the particles were considered to be delocalized. The particle analysis tool in Fiji was used for this quantification. Graphs were assembled using Graphpad Prism9.

---

**Supplemental Figure 1.**
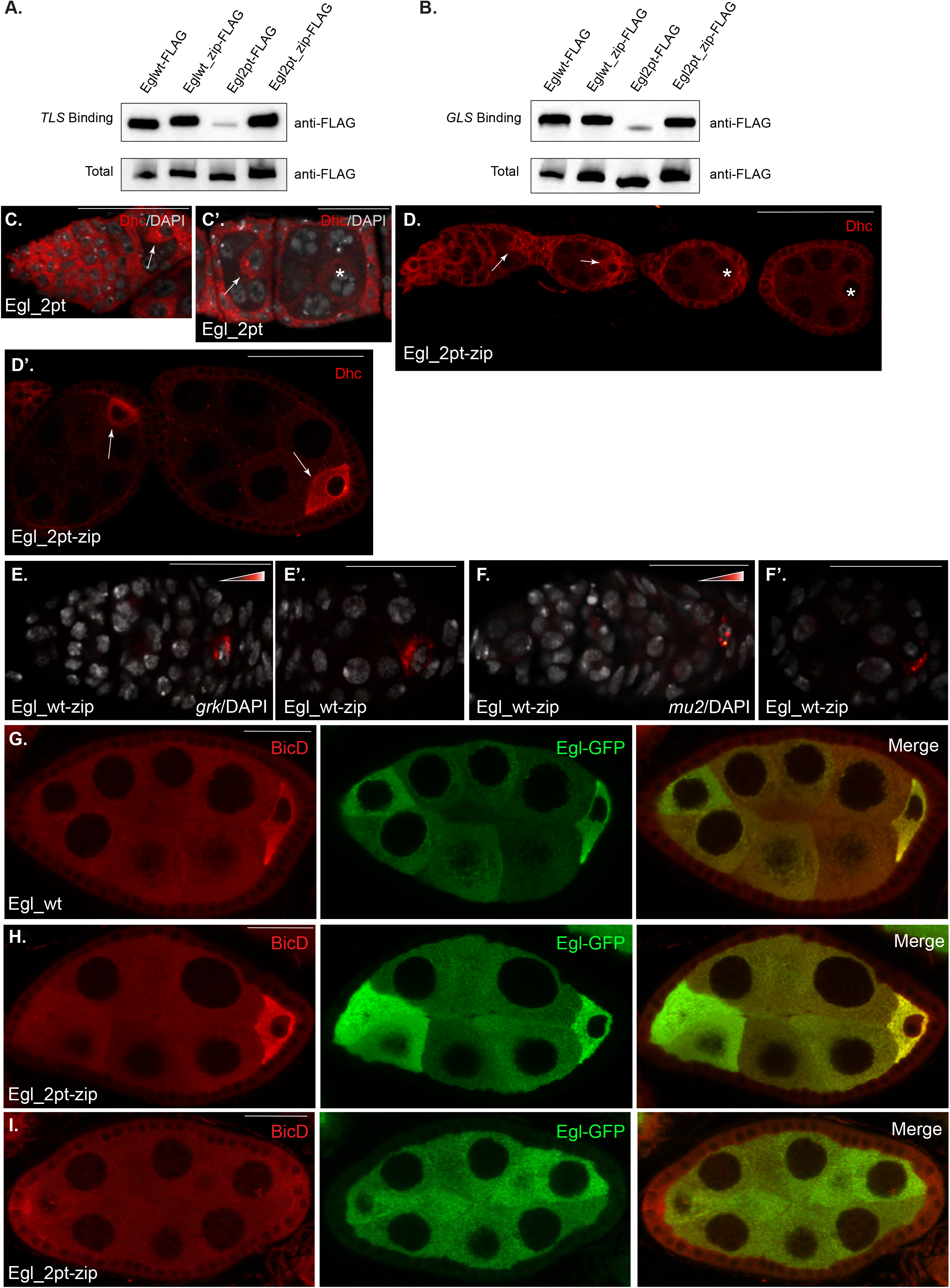
**(A-B)** Ovarian lysates from the indicated genotypes were prepared and used in an in vitro RNA binding experiment using the *TLS* (A) or *GLS* (B) localization elements. The bound proteins were eluted and analyzed by western blotting using the FLAG antibody. Total and bound fractions are shown. **(C-D)** Localization of Dynein heavy chain (Dhc, red) in strains expressing Egl_2pt (C, C’) or Egl_2pt-zip (D, D’). The egg chambers in panel C were also counterstained with DAPI (shown in greyscale). The enrichment of Dhc within the oocyte is indicated by the arrow. The asterisk indicated egg chambers that have lost oocyte fate. **(E-F)** Egg chambers from Egl_wt-zip are shown. The egg chambers were processed for smFISH using probes against *grk* (E, E’) or *mu2* (F, F’). The scale bar in D’ is 50 microns. In the rest of the images, the scale bar is 20 microns. **(G-I)** Egg chambers from Egl_wt and Egl_2pt-zip were processed for immunofluorescence using antibodies against BicD and GFP. In egg chambers expressing Egl_wt, both Egl-GFP and BicD were enriched within the oocyte (G). When oocyte fate was retained in Egl_2pt-zip flies, Egl-GFP and BicD were enriched within the oocyte (H). However, when oocyte fate was lost, neither protein was enriched within a single cell (I).

